# Gait speed is a biomarker of cancer-associated cachexia decline and recovery

**DOI:** 10.1101/2023.11.13.566852

**Authors:** Ishan Roy, Ben Binder-Markey, Danielle Sychowski, Amber Willbanks, Tenisha Phipps, Donna McAllister, Akash Bhakta, Emily Marquez, Dominic D’Andrea, Colin Franz, Rajeswari Pichika, Michael B. Dwinell, Prakash Jayabalan, Richard L. Lieber

## Abstract

**Background:** Progressive functional decline is a key element of cancer-associated cachexia. No therapies have successfully translated to the clinic due to an inability to measure and improve physical function in cachectic patients. Major barriers to translating pre-clinical therapies to the clinic include lack of cancer models that accurately mimic functional decline and use of non-specific outcome measures of function, like grip strength. New approaches are needed to investigate cachexia-related function at both the basic and clinical science levels.

**Methods:** Survival extension studies were performed by testing multiple cell lines, dilutions, and vehicle-types in orthotopic implantation of K-ras^LSL.G12D/+^; Trp53^R172H/+^; Pdx-1-Cre (KPC) derived cells. 128 animals in this new model were then assessed for muscle wasting, inflammation, and functional decline using a battery of biochemical, physiologic, and behavioral techniques. In parallel, we analyzed a 156-subject cohort of cancer patients with a range of cachexia severity, and who required rehabilitation, to determine the relationship between gait speed via six-minute walk test (6MWT), grip strength (hGS), and functional independence measures (FIM). Cachectic patients were identified using the Weight Loss Grading Scale (WLGS), Fearon consensus criteria, and the Prognostic Nutritional Index (PNI).

**Results:** Using a 100-cell dose of DT10022 KPC cells, we extended the survival of the KPC orthotopic model to 8-9 weeks post-implantation compared to higher doses used (p<0.001). In this Low-dose Orthotopic (LO) model, both progressive skeletal and cardiac muscle wasting were detected in parallel to systemic inflammation; skeletal muscle atrophy at the fiber level was detected as early as 3 weeks post-implantation compared to controls (p<0.001). Gait speed in LO animals declined as early 2 week post-implantation whereas grip strength change was a late event and related to end of life. Principle component analysis (PCA) revealed distinct cachectic and non-cachectic animal populations, which we leveraged to show that gait speed decline was specific to cachexia (p<0.01) while grip strength decline was not (p=0.19). These data paralleled our observations in cancer patients with cachexia who required rehabilitation. In cachectic patients (identified by WLGS, Fearon criteria, or PNI, change in 6MWT correlated with motor FIM score changes while hGS did not (r^2^=0.18, p<0.001). This relationship between 6MWT and FIM in cachectic patients was further confirmed through multivariate regression (r^2^=0.30, p<0.001) controlling for age and cancer burden.

**Conclusion:** Outcome measures linked to gait are better associated with cachexia related function and preferred for future pre-clinical and clinical cachexia studies.

## Introduction

Cachexia is a muscle wasting syndrome that accounts for 20% of cancer mortalities and causes disability in 50-75% of cancer patients [1,2]. In addition to significant weight loss, a hallmark of cachexia is progressive functional decline, leading to decreased quality of life and limited tolerance to future cancer therapies[1]. While several treatments developed for cachexia successfully attenuate weight loss, none have been approved by the FDA due to their inability to improve physical function[3]. A major limitation of prior clinical trials is a lack of agreement on the outcome measure that best reflects changes in physical function during cachexia. Without identifying the best tools to measure function specific to cachexia, function-focused cachexia treatments cannot successfully be deployed [4]. Current clinical cachexia studies rely upon grip strength as the primary measure of physical function, as it correlates well with mortality in cancer and aging [5]. However, the role of grip strength in predicting function during cachexia, as opposed to mortality, is unclear[6]. No proxy measure of function from a cachexia clinical trial has been directly compared to measures of functional independence, a major indicator of quality of life[7,8].

Unfortunately, the situation is even more dire in pre-clinical cachexia studies. Though numerous functional measures are currently in use [9], there have been no attempts to define the relationship among these measures and cachexia. Additionally, pre-clinical investigations are limited to animal models of cachexia that do not reflect changes in human physical function, namely, the slow decline in function associated with disease progression. While existing animal cachexia models have yielded a plethora of molecular mechanisms and potential pharmaceutical targets, these models either succumb to tumor burden too quickly[10,11], or express genotypically or phenotypically uncommon cancers in humans [12,13]. As a result, these models are limited in their translation to either human cancer or physical function. New approaches to pre-clinical modeling of cancer cachexia are needed before development of translational treatments targeting function can occur [3].

This report addresses both the need to develop a new pre-clinical model of cachexia that leads to progressive functional decline and the need to define functional outcomes that bridge the pre-clinical and clinical settings. The first aim of this study was to extend the survival and cachexia windows of an orthotopic and syngeneic pre-clinical cachexia model, while preserving a human cancer phenotype, so that it may display functional decline and a wider therapeutic window for future interventions. The second aim was to validate our pre-clinical findings in a broader human cancer cachexia cohort in an inpatient rehabilitation facility (IRF), where high resolution function metrics are routinely collected. Using these parallel animal and human experimental systems, we show, that gait changes are the best predictors of functional outcome in cachexia and thus should be the outcome measure of choice for future clinical and pre-clinical studies.

## Methods

### Cachexia animal model

Animals were housed in 12-hour dark/light cycles with free access to standard rodent chow and water. Male C57BL/6J mice were obtained from Jackson laboratory (Bar Harbor, ME, USA). At 12 weeks old, mice underwent pancreatic orthotopic implantation with cells or sham injection using our prior techniques [14,15]. Injection vehicles included 100 µL sterile phosphate buffered saline (PBS) or Matrigel (Corning - Glendale, Arizona, USA) diluted 1:4 in PBS. Cell lines included FC1199, FC1242, and DT10022 derived from K-ras^LSL.G12D/+^; Trp53^R172H/+^; Pdx-1-Cre (KPC) mice [15]. Post-operatively, animal pain and suffering were monitored through daily body condition scoring (BCS) in accordance with IACUC guidelines, and were given meloxicam and Buprenorphine SR for the first 48 hours. No animals exhibited significant pain behaviors via the BCS related to the surgery after 48 hours until 2 weeks, when wound clips were removed. In “survival” studies, animals were euthanized after meeting IACUC morbidity criteria. In serial studies, animals were euthanized on a weekly basis.

### Ex vivo tissue and histologic processing

Mouse tissues were frozen in liquid nitrogen-cooled isopentane or fixed in 10% formaldehyde [16]. Prior to freezing/fixation, individual wet weight of all muscle was obtained. TA muscle sections were stained for laminin and cross sectional area was calculated using Myovision 1.0 (see supplemental methods). Pancreatic tissue was stained with hematoxylin and eosin (H&E), Alcian Blue, and Masson’s Trichrome (MTC). Liver and lung sections were stained with H&E or cytokeratin-19 (DSHB TROMA-III).

### Biochemical analysis

Hydroxyproline was measured using our prior approach [17]. Each hydrolyzed sample was evaporated along with L-hydroxyproline standards (2-50 ug/ml) in a desiccator followed by color development using p-diamino-benzaldehyde and Chloramine-T. The absorbance was read at 550nm. Hydroxyproline content was converted to collagen by multiplication factor 7.46.

### Animal behavior analysis

Mice underwent a 3-day acclimatization period for functional assay environments. A grip strength meter (Columbus Instruments) was used to measure the grip strength of each individual limb, with hindlimb grip strength calculated as an average of both left and right limbs (see Supplementary Methods). For open field testing, the measurements of the chambers were 56cm x 56cm. Mice were placed in the center of the chamber and allowed to explore the chamber for 5 minutes. The center, or zone 2, was defined as a 3 x 3 grid within the larger 5 x 5 space. For Y-maze, each arm of the maze was 60 cm in length, 3.5 cm wide at the bottom, and 14 cm wide at the top. Animals were placed in the middle arm of the maze and allowed to explore the maze undisturbed for 8 mins. All movements were recorded using LimeLight4 software (Actimetrics) and average gait speed was calculated as previously reported [18].

### Human subject study design

This was a retrospective study of cancer patients admitted to a free-standing inpatient rehabilitation facility (IRF) from March 2017 to September 2018 [19]. Inclusion criteria were: ≥18 years of age, prior oncologic care at an affiliated comprehensive cancer center, and primary rehabilitation impairment due to cancer or its treatment. Any patients without available weight/height data, or six-minute walk test (6MWT)/hand grip strength (hGS) performed at admission and discharge from rehabilitation were excluded.

250 patients matched initial inclusion criteria. 28 patients were excluded for cancer care at a non-affiliated institution and 66 patients were excluded for incomplete data for a total of 156 patients included. As required by Centers for Medicare and Medicaid Services (CMS), all patients received a minimum of 900 minutes per week of a combination of physical (PT), occupational (OT), and speech and language pathology (SLP) therapies. Electronic and manually extracted data included details of patient’s acute care, rehabilitation care, and cancer courses up to 6 months before and after their IRF admission.

### Human cachexia cohort selection

The Weight Loss Grading Scale (WLGS) [20] was used for identifying patients with varying cachexia severity [21]. Secondary cachexia markers were consensus weight-based criteria (5% weight loss in 6 months or 2% weight loss in 6 months with BMI< 20) [22], and Prognostic Nutritional Index (Albumin level (g/L) + 0.005 × Lymphocyte count per microliter) (PNI), shown in several studies of cancer patients to be linked to malnutrition, cachexia, and negative clinical outcomes [23]. The threshold for PNI was set at ≤40, or serious malnutrition [24].

### Human functional outcome measures

All patients had admission and discharge Functional Independence Measure (FIM) scores, the gold standard measure of function at IRFs [25], and an 18-item scale (scored from one to seven) separated into 13 motor item and 5 cognitive subsets. Change in FIM scores from admission to discharge were the primary functional measures used, with 6MWT and hGS also captured at admission and discharge. All measures were obtained by certified rehabilitation therapists as part of standard care.

### Statistical analysis

Significance (α) was set to 0.05 for all analyses. Analyses were completed in Graphpad Prism 9 and IBM SPSS Statistics 26. Time-to-event analyses were done by Kaplan-Meier. Simple linear regression was used for correlations. Pair-wise analyses were performed using Student’s T-test. Multiple cohort analysis at single times were performed by one-way ANOVA. Two-way ANOVA analysis was performed for multiple cohorts over time, with Tukey *post-hoc* analysis. ANCOVA was performed to generate body-weight adjusted muscle mass and muscle mass adjusted collagen content means. For human subjects, descriptive data were analyzed by chi-squared analysis. Binary relationships were analyzed by logistic regression. Multivariate linear regression included FIM gain as the dependent variable and age, disease recurrence status, 6MWT, and hGS as co-variates. All values in figures are mean ± SEM unless otherwise indicated. See supplemental methods for principal component analysis details.

## Results

### Cell titration extends survival in orthotopic pancreatic cancer mouse model while preserving human cancer phenotype

Current commonly used cachexia models succumb to tumor burden in 3-4 weeks, a timeline that is too short to quantify longitudinal functional change[9]. The orthotopic allograft KPC model has emerged as a preferred, syngeneic pre-clinical model for cachexia given that it genotypically and phenotypically models human pancreatic cancer[26], but as currently used also has a limited survival window of only 3-4 weeks. To prolong survival in this model, we tested a variety of KPC cell lines, cell doses, and injection vehicles (Fig. 1A). Amongst several KPC cells lines [15], the 10022 cell line was selected, as it had significantly longer median survival (p<0.001) (Supplemental Fig. 1A). We then performed a dose-titration experiment with 10022 cells over five orders of magnitude ranging from 10 to 1 million cells. The dose of 100 cells extended survival to greater than 8 weeks (p<0.001) (Fig. 1B) while reliably inducing a tumor. *Ex vivo* analysis of pancreatic and primary tumor mass revealed that both the lowest (r^2^ = 0.31, p<0.001) and highest (r^2^ = 0.73, p<0.001) doses of 10022 cells maintained significant correlation between tumor size and time, as did intermediate doses (Fig. 1C, Supplemental Fig. 1C). We also compared PBS and Matrigel as injection vehicles and found no significant difference in survival between substrates (Supplemental Fig. 1B). Consistent with previous KPC models, we observed histologic hallmarks of metastatic pancreatic cancer using the 100-cell dose. Primary tumors revealed ductal adenocarcinoma morphology on H&E, increased glycosaminoglycan secretion via Alcian blue staining, and increased collagen deposition in the tumor ECM via Masson’s trichrome (Fig. 1D, yellow brackets), all of which are structural features of human pancreatic cancer[27]. Using gross dissection identification, CK19, and H&E staining we identified pancreatic cells metastasized to the peritoneum, liver, and lung within the 100-cell dose (Fig. 1D), median time to all types of metastasis was 7 weeks (Supplemental Figs. 1D-E). Thus, 100 cells of the 10022 cell line, when orthotopically implanted, lead to an expanded survival window of 8-9 weeks while preserving a human cancer phenotype.

**Fig. 1.**
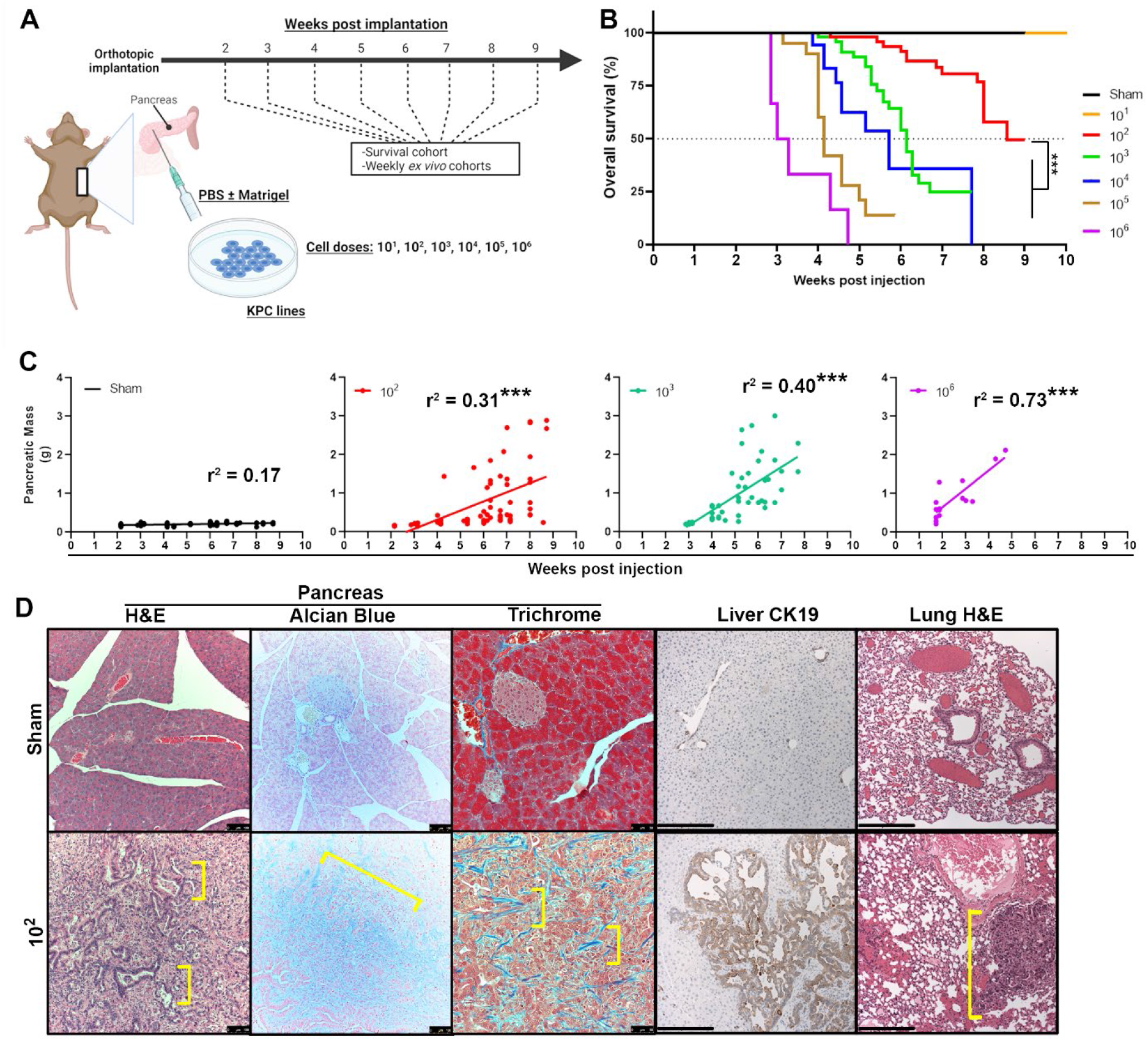
Serial titration of KPC cells generates an orthotopic implantation model with extended survival, progressive pancreatic tumor growth, and metastatic disease. (A) Study design. In a series of experiments, mice underwent pancreatic orthotopic implantation of KPC cells with the goal of evaluating distinct cell lines, cell doses (range: 10 to 1,000,000), and injection vehicles for extending survival of tumor-implanted animals. Animals were euthanized for *ex vivo* analysis either in weekly cohorts starting 2 weeks post-implantation, or at time of morbidity (survival), in accordance with institutional guidelines. (B) Kaplan-Meier survival analysis of cohorts implanted with serial dilution of 10022 KPC cells. (D) Representative pancreatic tissue sections of 100 cells implanted and sham animals, including pancreas H&E (size bar = 100 µm), pancreas Alcian Blue (cyan = glycosaminoglycans) (size bar = 100 µm), pancreas Masson’s Trichrome (blue = collagen) (size bar = 50 µm), CK19 stained liver (size bar = 250 µm), and lung H&E (size bar = 250 µm). For panels B-E, n = 33 animal subjects for PBS, n = 51-52 for 100 and 1,000, n = 16 for 1,000,000, and n = 18-20 for 10,000 and 100,000. *** denotes p<0.001.

### Orthotopic implantation of 100 pancreatic cancer cells leads to skeletal and cardiac muscle wasting, increased collagen deposition, and systemic inflammation

To quantify cachexia syndrome within this new model, we first measured animal body-weight. At week 5 post-implantation, body-weight was 5% lower in the low-dose animals compared to controls (p<0.05) (Fig. 2A); though by week 7, this difference was confounded by tumor burden and peritoneal fluid accumulation, a common feature of pancreatic disease [27]. In pilot imaging studies *in vivo*, significant hindlimb muscle wasting was observed in 100-cell implanted mice by 6-7 weeks using both CT (p<0.01) and MRI (p<0.01) (Supplemental Fig. 2). Using whole muscle wet-weights, and controlling for initial animal body weight via ANCOVA, both total hindlimb skeletal muscle mass and cardiac muscle mass in 100-cell mice were significantly lower compared to sham mice (i.e., a significant dose x time interaction, p<0.0001) (Fig. 2B-C). Within the 100-cell cohort, significant muscle weight decrease began 5 weeks post-implantation (Tukey p<0.05) (Fig. 2B-C). Individual hindlimb skeletal muscles in tumor mice (quadriceps, gastrocnemius, and tibialis anterior) demonstrated significant muscle weight decrease over time compared to sham animals (dose x time interaction: p<0.001, p<0.05, p<0.01, respectively for each muscle), while hamstrings (semimembranosus) did not (Fig. 2D).

**Fig. 2.**
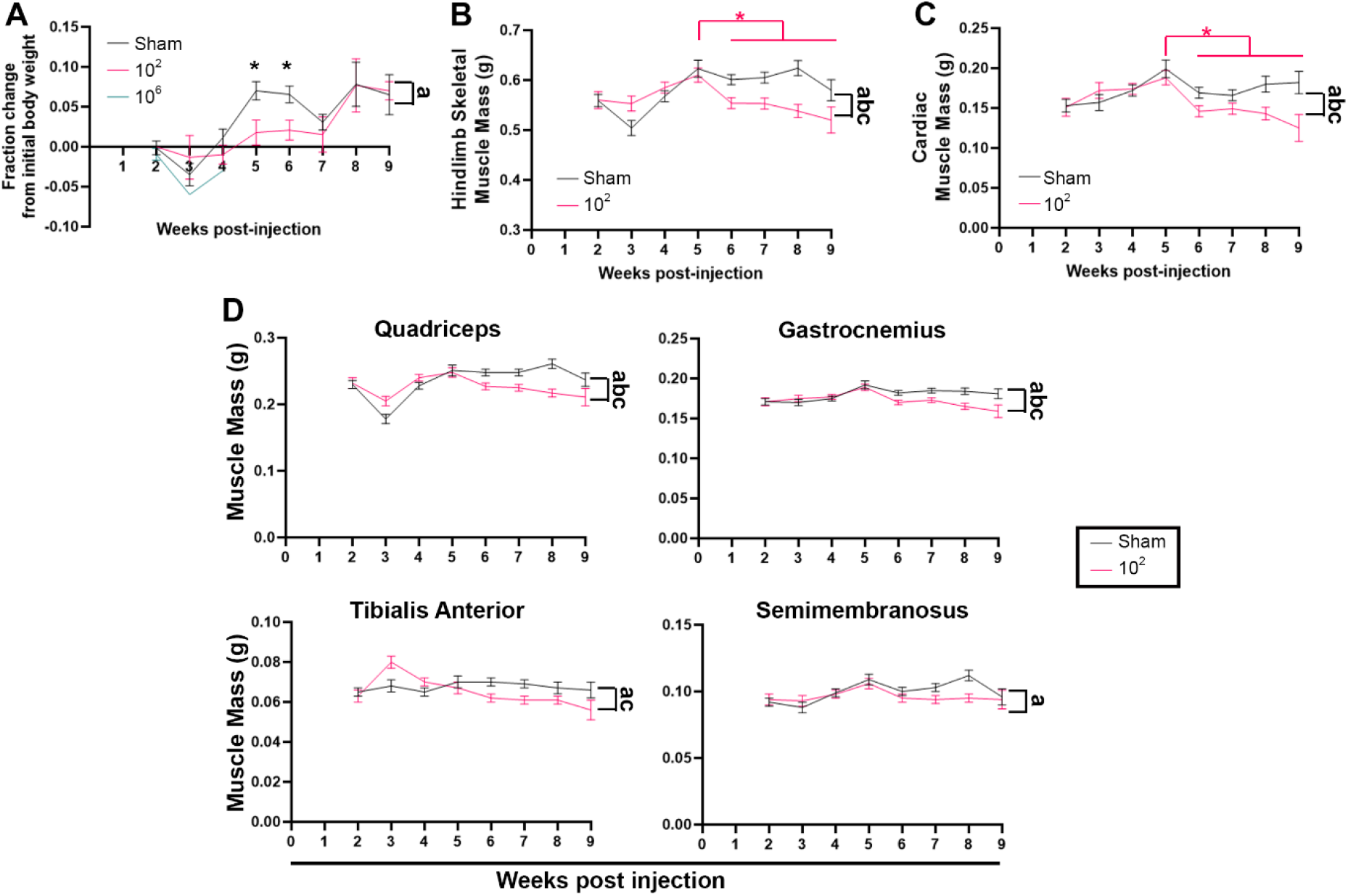
Low-dose KPC orthotopic model extends the window of muscle wasting. (A) Fraction change from pre-implantation body weight to final body weight in serial weekly PBS (sham), 100, and 1,000,000 cohorts. (B) Hindlimb skeletal muscle mass (sum of quadriceps, gastrocnemius, tibialis anterior, and semimembranosus wet weights) over serial weekly cohorts in sham and 100-cell implanted mice. (C) Cardiac mass over serial weekly cohorts in sham and 100 implanted animals. (D) *Ex vivo* mass of individual hindlimb muscles over time in serial weekly cohorts. For panels A-D, n = 7-12 animal subjects per weekly cohort per cell dose, other than week 9 (n=5 per cell dose). Black * indicates p < 0.05 in Tukey *post-hoc* analysis between cell doses. For the two-way ANOVA, “a” indicates p < 0.05 for time factor, “b” indicates p < 0.05 for cell dose factor, and “c’” indicates p < 0.05 for a time x dose interaction. Pink * indicates p < 0.05 in Tukey *post-hoc* analysis between time points within the 100 cohort.

To define precisely the onset of muscle wasting, we measured tibialis anterior muscle fiber cross-sectional area via laminin staining. In weeks 3 through 8 post-implantation, 100-cell animals had significantly lower mean fiber area compared to sham mice (p<0.001) (Figs. 3A-B). Given that muscle fibrosis has been previously noted in cachexia [28], we tracked skeletal muscle collagen content over time and found that the collagen content increased in tumor mice compared to sham animals (dose x time interaction, p<0.01), with a significant within cohort difference observed by 6 weeks post implantation (Fig. 3C) (Tukey, p<0.05). To correct for fractional collagen content increase due to decreased fiber size, we also recalculated collagen content using tibialis anterior (TA) muscle mass as a covariate (ANCOVA), still found significantly increased collagen content (p<0.0001) (Fig. 3D), and noted increased perimysial collagen deposition in the TA muscles of tumor animals in Trichrome staining (Fig. 3E).

**Fig. 3.**
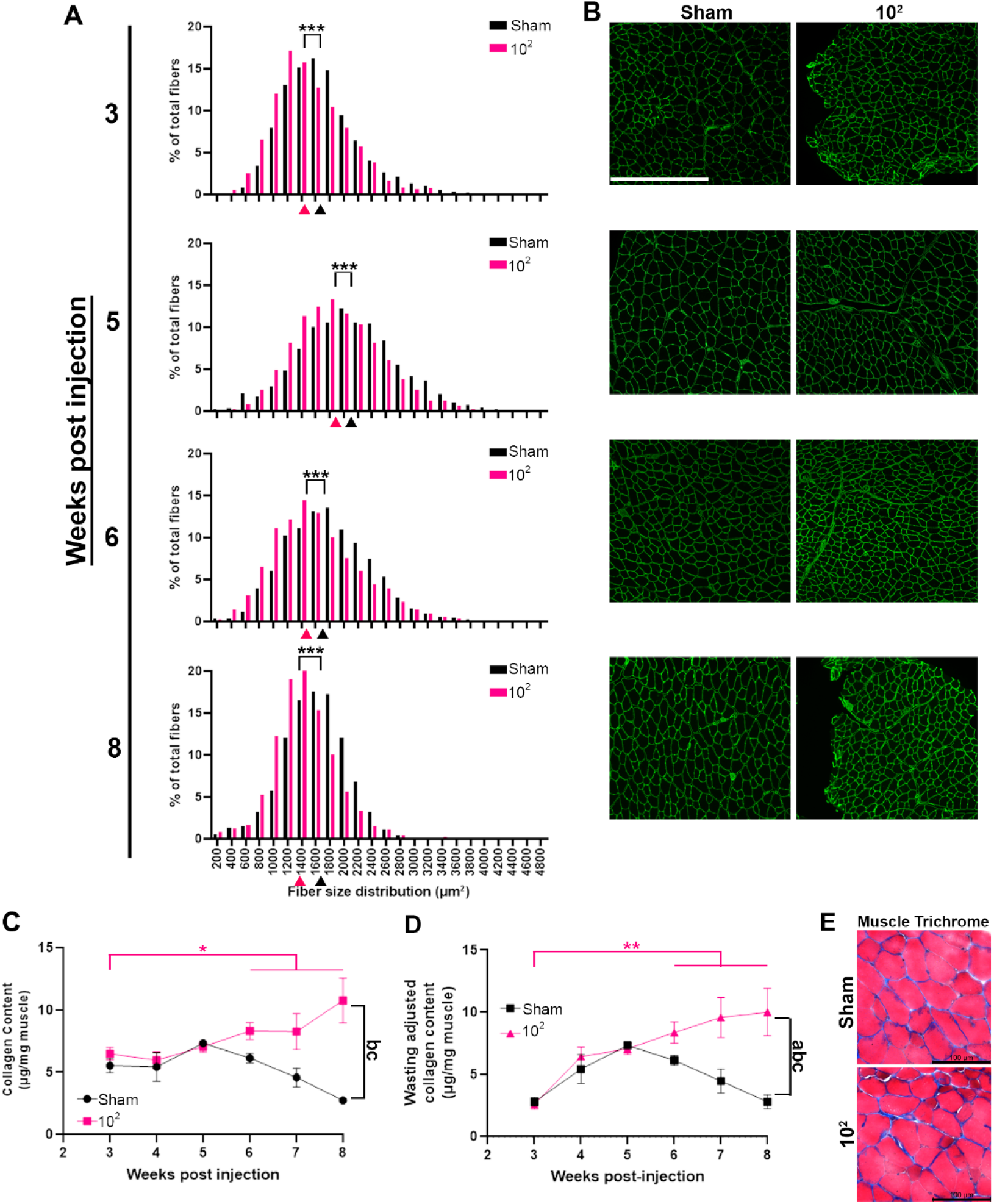
Skeletal muscle fiber size decreases and collagen content increases over time. (A) Histogram of tibialis anterior fiber cross-sectional area in PBS and 100 implanted mice in weekly cohorts. Colored triangles below abscissa indicate cohort means. *** indicates p < 0.001 in unpaired t-test. (B) Representative laminin labeled immunofluorescence images for each cell dose and time cohort. Scale bar size = 500 µm. (C) Tibialis anterior collagen content in weekly sham and 100 cohorts. n = 6-11 per time point (D) Tibialis anterior collagen content with means adjusted for muscle mass in weekly sham and 100 cohorts. (E) Representative Masson’s Trichome images indicating perimysial collagen localization in PBS and 100 animals at 8 weeks post-implantation. Scale bar size = 100 µm. For panels C-D, key as in Fig. 2.

Next, we characterized systemic inflammation and noted significant splenomegaly over time in 100-cell animals compared to controls (dose x time interaction, p<0.05), first occurring 4 weeks post implantation (Tukey p<0.01) (Supplemental Fig. 3A). We sampled germinal centers in a subset of spleens from sham and tumor animals and found that there was a trend towards expansion of the marginal zone area in tumor animals (p=0.09) (Fig. Supplemental Fig. 3B-C), indicating an inflammatory etiology for splenomegaly rather than vascular congestion. Using the mouse sepsis score (MSS), 100-cell animals also displayed significantly increased sepsis behavior over time compared to sham animals (dose x time interaction, p<0.0001), as early as 6 weeks post implantation (Supplemental Fig. 3D). At the blood level, we found significant serum Pentraxin-2 and GDF15 elevation (p<0.01) in tumor animals compared to sham, though without a time trend (Supplemental Fig. 3E).

Taken together, these physiologic data demonstrate that progressive muscle wasting occurs over time in the setting systemic inflammation in our low-dose model.

### Behavioral phenotyping shows time-dependent functional decline in low-dose cachexia mouse model

We next characterized physical function using a battery of rodent behavioral assays. We first examined grip strength as it is the most often used functional parameter in animal-based cachexia studies. We found there was a significant decline in hindlimb grip strength over time in 100-cell animals compared to sham (dose x time interaction, p<0.0001). However, *post-hoc* analysis of the tumor cohort yielded no threshold timepoint for sustained decline (Fig. 4A). Instead, when the results were plotted in reverse from time of mortality (Supplemental Fig. 4A), we found that grip strength in tumor animals only declined within 1 week of mortality, suggesting it may not be a measure of longitudinal changes. Notably, individual hind- and fore-limb grip strength strongly correlated with each other in 100-cell animals (Supplemental Fig. 4B), suggesting that individual and combined limb grip strengths do not have different sensitivity. As a measure of animal coordination, we also performed rotarod testing, which showed no significant difference between tumor and sham animals across time, albeit with a transient decline at 2 weeks post-implantation (Tukey, p<0.05) (Supplemental Fig. 4C).

**Fig. 4.**
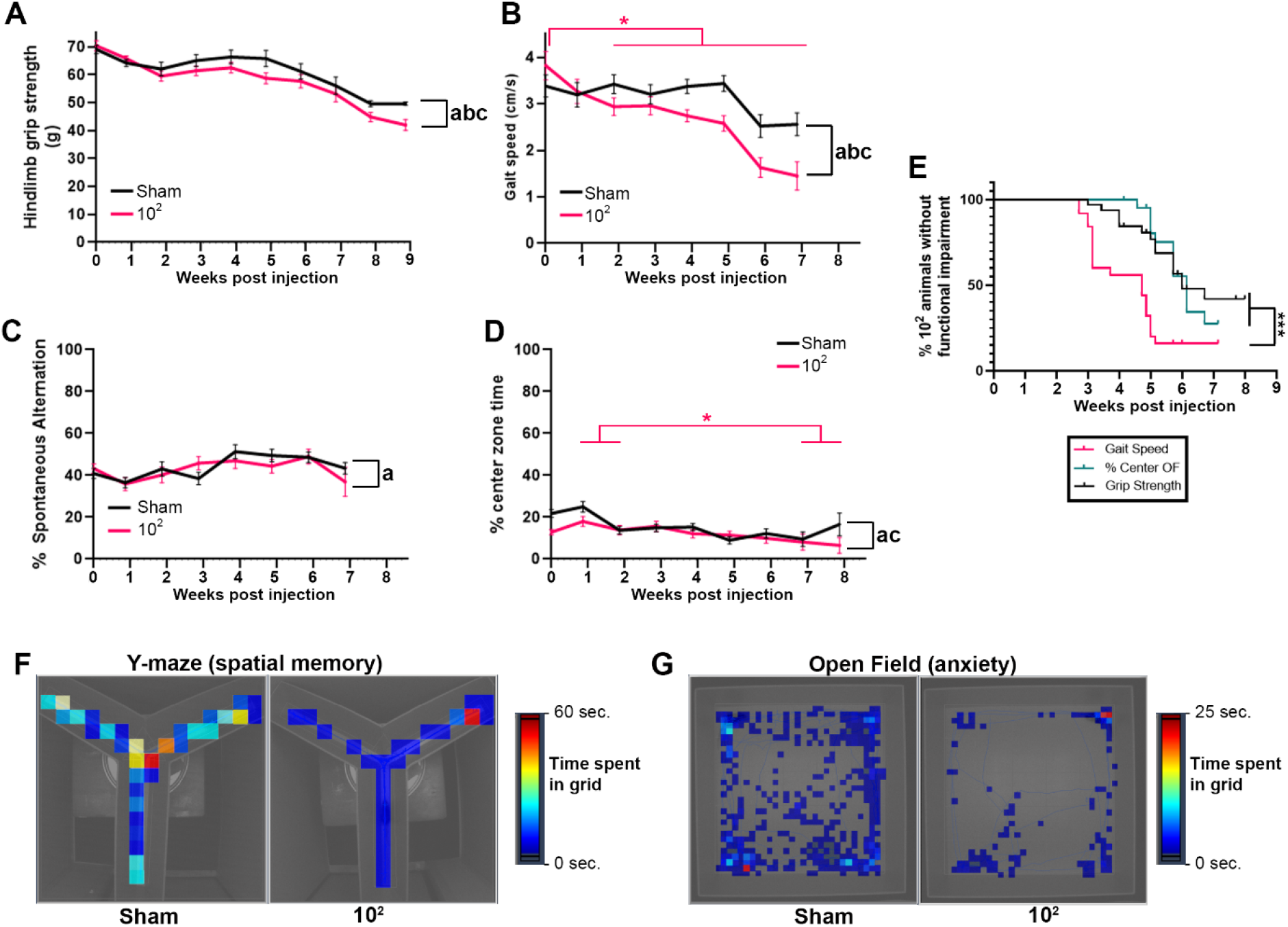
Gait speed decline is the earliest indicator of functional decline. (A) Hindlimb grip strength in sham and 100 cohorts vs. weeks post-implantation. (B) Self-selected linear path gait speed in the Y-maze in sham and 100 cohorts. Pink * indicates p < 0.05 in Tukey post-hoc analysis between time points within the 100 cohort. (C) Spontaneous alternation percentage between Y-maze arms in sham and 100 cohorts. (D) Percentage time spent in center zone of open field maze in sham and 100 cohorts. (E) Kaplan Meier time to event analysis for impairment in gait speed, grip strength, and % center zone time in 100 cohort animals. Impairment defined as below baseline value. (F&G) Representative heat maps of time spent in grid locations in Y- and open field mazes at 7 weeks post implantation. For panels A-G, n = 26-33 per cohort. Key as in Fig. 2.

To measure animal function more sensitively, we performed maze-based assays. Y-maze and open field assays were first designed to simultaneously estimate locomotor activity and cognitive behavior [18,29], allowing us to account for central behavior changes during physical functional decline due to cancer. In both the Y-maze and open field settings, self-selected straight line gait speed was significantly decreased over time in tumor mice compared to sham (dose x time interaction, p<0.01), with significant within tumor cohort decline from baseline occurring as early as 2 weeks post-implantation (Tukey, p<0.05) (Figs. 4B, 4F) (Supplemental Fig. 4D). We did not find evidence of cognitive decline in the cancer group as measured by spontaneous alternation within the Y-maze (Figs. 4C, 4F). Time spent in the center zone was significantly decreased over time within the tumor group (dose x time interaction, p<0.01), indicating increased anxiety[18], though this did not occur until 7-8 weeks post-implantation (Tukey p<0.05) (Figs. 4D, 4G). Finally, in a Kaplan-Meier analysis of all types of functional decline, median time to impairment in gait speed occurred 2 weeks earlier than impairment in grip strength or anxiety (p<0.0001).

In total, given the distinct longitudinal physiologic and functional phenotype of this model, we will now refer to this Low-dose Orthotopic model as the “LO” model.

### Multivariate analysis demonstrates that gait speed is specifically associated with cachexia and cancer, while grip strength is only associated with cancer

We noted heterogeneity over time in the physiologic data collected from LO animals and we hypothesized that, within the model, there may be distinct cachectic and non-cachectic populations. To address this, we performed principal component analysis (PCA) using 128 LO animals confirmed histologically to have primary tumor. To represent cancer, muscle wasting, and inflammation we included primary tumor size, quadriceps muscle mass, cardiac muscle mass, peritoneal metastasis, and splenomegaly as variables. PCA yielded two orthogonal principal components (PC) (Supplemental Table 1). PC1 correlated positively with primary tumor, splenomegaly, and metastasis and negatively with quadriceps and cardiac muscle while PC2 correlated positively with all five variables (Fig. 5A, Supplemental Table 1). Thus, PC1 represents cancer with cachexia, whereas PC2 represents cancer without cachexia. When individual subject PC scores were mapped over time, there was a positive shift toward PC1 (“cancer + cachexia”) over time, while a smaller cohort of animals at primarily earlier time points shifted toward PC2 (“cancer”) (Fig. 5B), which mirrors the parallel progression of cachexia alongside cancer seen in humans. Using the PCA, we then asked whether any functional measures (gait speed, grip strength, open field anxiety) were independently associated with cachexia versus cancer components via linear regression, and found that gait speed was the only functional measure independently associated with all five variables in the PCA, including muscle, tumor, and inflammation parameters (Table 1). In contrast, grip strength was only associated with tumor and inflammation and open field anxiety had no significant associations in the PCA (Table 1).

**Fig. 5.**
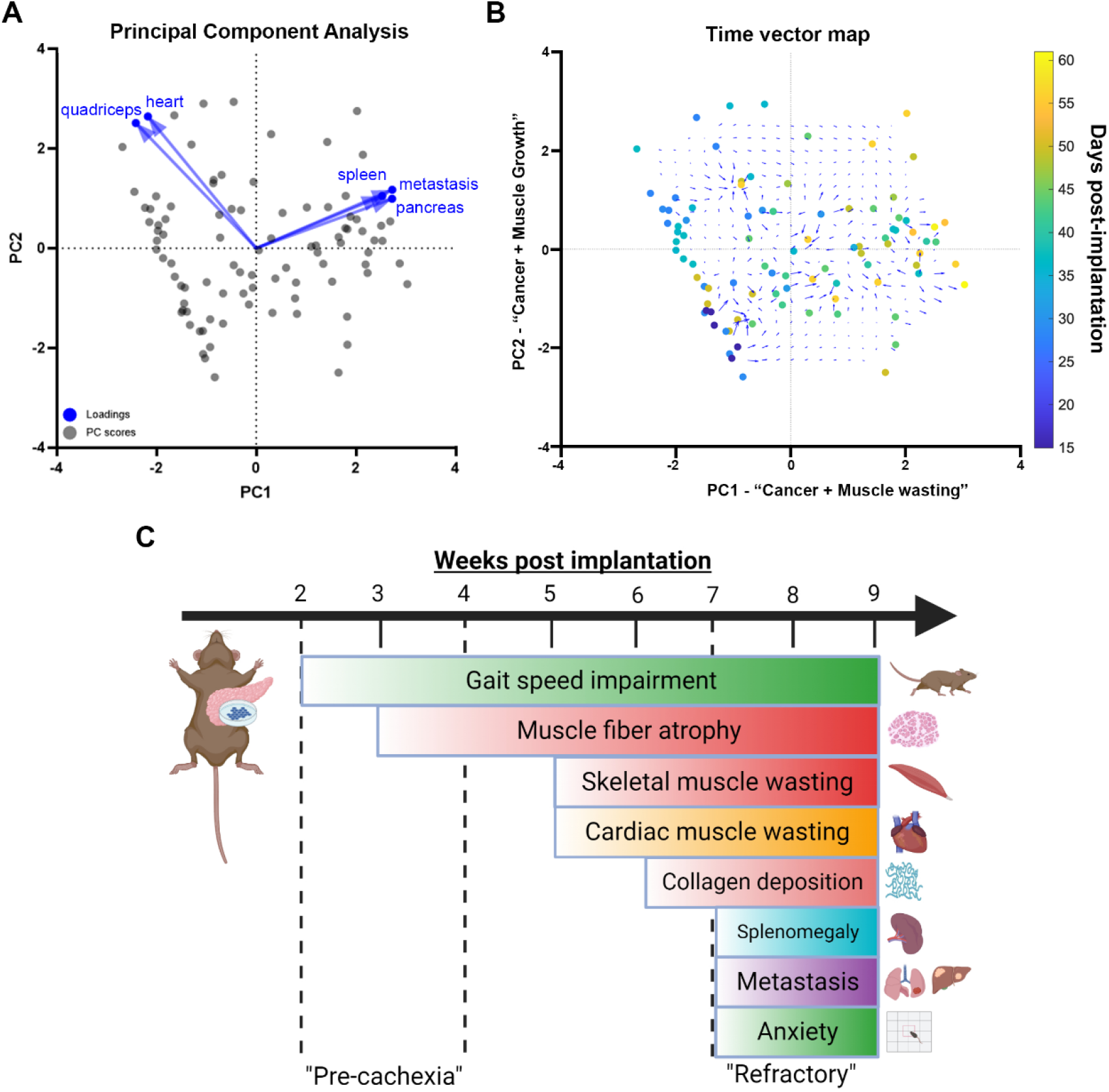
Principle component analysis (PCA) of pancreatic cancer model reveals distinct cancer cachectic and non-cachectic components stratified over time. (A) PCA results with individual PC2 *vs.* PC1 scores of all 128 low-dose animals included, along with loadings for five physiologic variables (pancreas mass, metastasis, spleen mass, quadriceps mass, and heart mass). (B) Two-dimensional PC2 *vs*.PC1 plot overlayed with days since injection (blue to yellow gradient, right) and contour mesh of time vectors (directions pointing towards increasing magnitude of time). (C) Timeline of physiologic (muscle = red/orange, systemic = purple, inflammation = blue) and functional (green) changes in low-dose KPC orthotopic implantation model. Proposed pre-cachexia and refractory cachexia phases are annotated with dashed lines.

**Table 1.** Principal component regression analysis of functional measures with physiology variables.

| **Functional measure** | **Variable** | **Estimate** | **Standard error** | **95% CI (asymptotic)** | **P value<sup>1</sup>** |
| --- | --- | --- | --- | --- | --- |
| **Gait Speed** | **quadriceps** | 12 | 3.5 | 4.5 to 20 | 0.0044* |
|  | **Pancreas** | -0.45 | 0.054 | -0.57 to -0.34 | <0.0001* |
|  | **Heart** | 9 | 2.9 | 2.8 to 15 | 0.0076* |
|  | **Spleen** | -5 | 0.63 | -6.4 to -3.7 | <0.0001* |
|  | **peritoneal metastasis** | -0.7 | 0.086 | -0.88 to -0.51 | <0.0001* |
| **Grip Strength** | **quadriceps** | -56 | 41 | -142 to 30 | 0.19 |
|  | **Pancreas** | -5.9 | 0.62 | -7.2 to -4.6 | <0.0001* |
|  | **Heart** | -57 | 34 | -127 to 13 | 0.10 |
|  | **Spleen** | -71 | 7 | -85 to -56 | <0.0001* |
|  | **peritoneal metastasis** | -9.6 | 0.96 | -12 to -7.6 | <0.0001* |
| **% Center** | **quadriceps** | 14 | 78 | -155 to 183 | 0.86 |
|  | **Pancreas** | 0.56 | 0.95 | -1.5 to 2.6 | 0.56 |
|  | **Heart** | 12 | 63 | -126 to 151 | 0.85 |
|  | **Spleen** | 6.9 | 11 | -17 to 31 | 0.55 |
|  | **peritoneal metastasis** | 0.93 | 1.5 | -2.3 to 4.2 | 0.55 |

1. See Methods for principle component analysis details.

Thus, using the LO model, we were able to delineate the specific timing of a breadth of physiologic, biochemical, and functional events in relation to both cancer and cachexia (Fig. 5C).

### In human cancer patients with cachexia, gait speed, but not grip strength, correlates with improvements in motor function during rehabilitation

Given the finding that gait speed is the better functional indicator of cachexia in the LO model, we hypothesized that gait impairments may also be superior to grip strength to detect functional change in human cancer patients. To address this, we analyzed a cohort of 156 patients admitted to our IRF for cancer-related functional decline (Supplemental Table 2). Each patient had functional independence measure (FIM) scores, handgrip strength (hGS) and six-minute walk test distance (6MWT) collected at admission and discharge (Fig. 6A). The cohort demonstrated significant improvements in motor function, cognitive function, hGS, and 6MWT measurements during rehabilitation (Fig. 6B). Cachexia was highly prevalent in the 156-patient cohort (Fig. 6C). 121 patients had some degree of cachexia as estimated by the weight loss grading scale (WLGS)[20,21], 90 patients met Fearon *et al.* weight/BMI cachexia criteria[22], and 48 patients out of 76 patients with available serum met the blood-based Prognostic Nutritional Index (PNI) threshold for serious malnutrition (Fig. 6C) [23,24]. Of the baseline variables collected, only age was significantly associated with cachexia status (p<0.01) (Supplementary Table 2). Using univariate linear regression analysis, we found no correlation between hGS and motor FIM for cancer patients with any of the three cachexia identifiers (Fig. 6D). In contrast, there was a significant correlation between change in 6MWT and FIM motor change in both cancer patients with and without cachexia measured by each marker (Fig. 6E) (Supplementary Table 3), and that held true in both reported genders (Supplementary Table 4). In regression analysis between 6MWT and FIM cognitive changes, there was no significant correlation across all WLGS scores (Supplementary Table 5), indicating that cognition was not a major confounder.

**Fig. 6.**
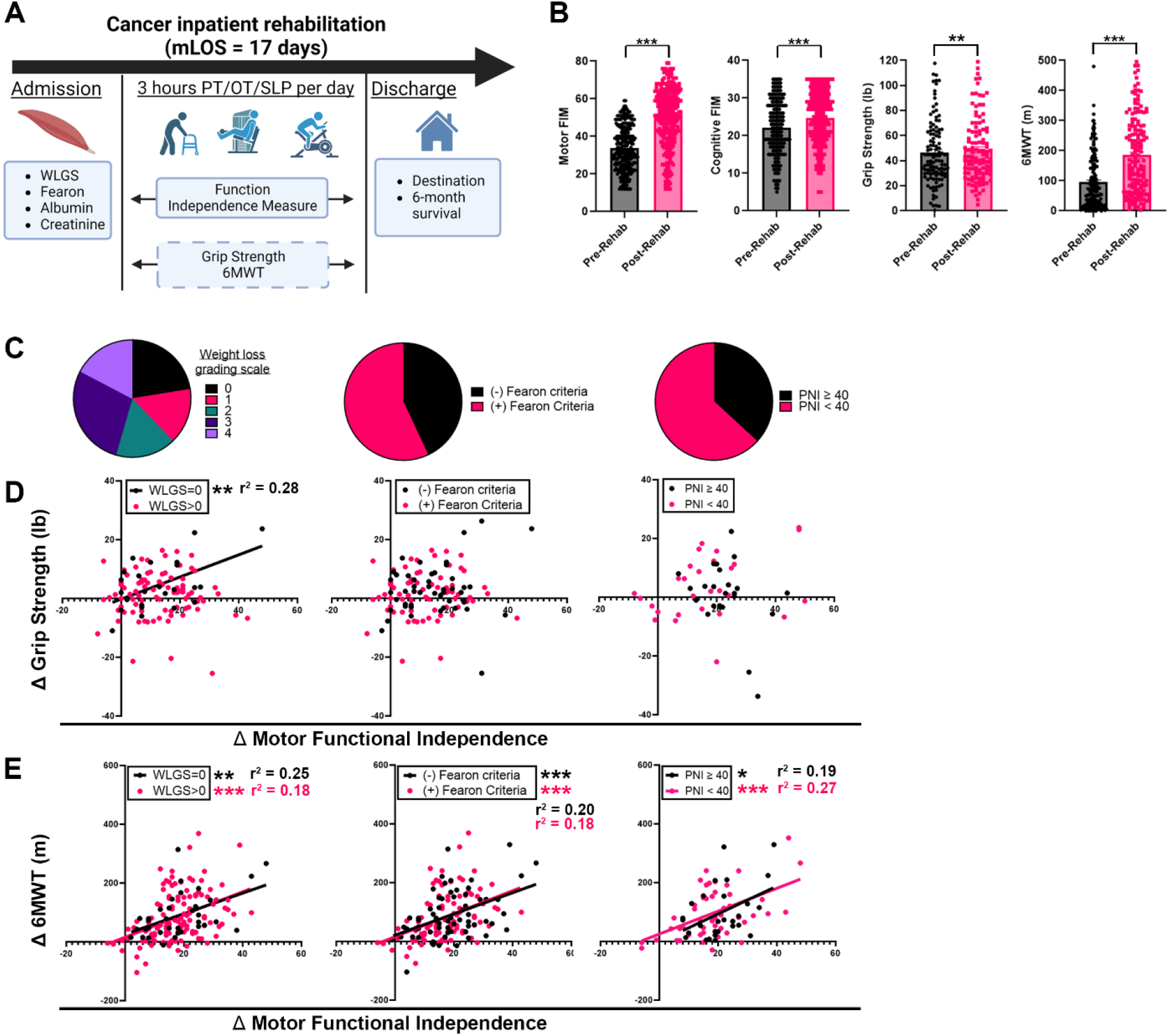
Gait speed, but not grip strength changes correlate with changes in motor functional independence in cancer patients with cachexia receiving inpatient rehabilitation. (A) Study Design. 156 patients with cancer who were admitted to our inpatient rehabilitation facility were retrospectively assessed for cachexia markers, including Weight Loss Grading Scale (WLGS), Fearon *et al.* criteria, Prognostic Nutritional Index (PNI). During inpatient rehabilitation, each patient received at least a total of 3 hours per day of physical therapy (PT), occupational therapy (OT), and speech/language therapy (SLP). Change in patients’ functional independence measure (FIM) motor and cognitive sub-scores at admission and discharge from rehabilitation were correlated with hand grip strength and six-minute walk test (6MWT) distance ambulated. Additional discharge measures included discharge destination and six-month mortality. (B) Motor FIM, cognitive FIM, grip strength, and 6MWT values pre- and post-rehabilitation. * indicates p < 0.05 in Mann-Whitney comparison. (C) Pie chart distribution of patients with muscle wasting using WLGS, Fearon *et al.* criteria, and severe PNI (black = no cachexia). (D-E) Scatter plots of change in grip strength or 6MWT vs. change in motor FIM in IPR cohort stratified by individual cachexia markers. * indicates p < 0.05 for linear regression slope significantly non-zero, with corresponding r^2^ goodness of fit.

In multivariate analysis to determine the influence of age and disease recurrence on functional observations, in cachectic patients with a WLGS > 0, we found 6MWT change independently correlated with FIM motor change, while hGS gain did not (Table 2). There were no independent associations with cognitive changes. Discharge destination (i.e., whether someone is deemed at a functional level able to remain independent in their home) is a major indicator of quality of life and in cachectic patients with WLGS > 0, we found significantly increased odds of discharge to a higher functional independence setting with increasing 6MWT walk test gains during their IRF stay (p<0.05) (Supplementary Table 6). In contrast, there was no association between hGS and discharge setting (p=0.60). In a multivariate analysis including age and disease recurrence, 6MWT change also trended toward an independent association with discharge setting (p=0.08).

**Table 2.** Multivariate analysis of cancer patients with Weight Loss Grading Scale > 0 and functional independence measures and selected clinical variables.

| WLGS<sup>1</sup>>0 | Estimate | Standard error | 95% CI (asymptotic) | P value<sup>4</sup> | R squared |
| --- | --- | --- | --- | --- | --- |
| **FIM<sup>2</sup> Motor gain** |  |  |  |  |  |
| **Δ 6MWT<sup>3</sup> (m)** | 0.064 | 0.014 | 0.035 to 0.092 | <0.0001 | 0.30 |
| **Δ Grip strength (kg)** | -0.19 | 0.17 | -0.54 to 0.14 | 0.26 |  |
| **Age** | 0.089 | 0.10 | -0.11 to 0.29 | 0.38 |  |
| **Recurrent disease** | -1.07 | 3.07 | -7.2 to 5.1 | 0.73 |  |
| **FIM Cognitive Gain** |  |  |  |  |  |
| **Δ 6MWT (m)** | 0.0070 | 0.0063 | -0.0055 to 0.019 | 0.27 | 0.033 |
| **Δ Grip strength (kg)** | -0.041 | 0.075 | -0.19 to 0.11 | 0.59 |  |
| **Age** | 0.026 | 0.044 | -0.062 to 0.11 | 0.55 |  |
| **Recurrent disease** | -0.50 | 1.34 | -3.2 to 2.1 | 0.71 |  |

1. WLGS = Weight loss grading scale 2. FIM = Functional Independence Measure 3. 6MWT = Six-minute walk test 4. Multivariate linear regression

These clinical findings parallel our pre-clinical data demonstrating that outcomes associated with gait are better indicators of physical function in cancer associated cachexia and thus should be considered the preferred outcome measures for monitoring cachexia related function.

## Discussion

The primary objective of this study was to identify a translational functional outcome measure for cancer-associated cachexia. We first developed an extended animal model that allowed us to determine the precise timing of physiologic and behavioral events during the progression of both cancer and cachexia. We now refer to this <u>L</u>ow-dose <u>O</u>rthotopic model as the “LO” model. We found that decline in gait speed was both an early event in the development of the LO model and specifically decreased in animals with cancer-associated cachexia. In contrast, grip strength decline developed much later in the model and was only linked to cancer progression and mortality, and not cachexia. In human subjects, we discovered a similar relationship in that 6MWT change correlated with functional independence in cancer patients with cachexia while handgrip strength change did not. Taken together, these data demonstrate that gait speed is a sensitive functional outcome measure for cancer cachexia in both clinical and pre-clinical settings and should be strongly considered for inclusion in future translational and clinical studies.

Both LLC and C26 models were commonly used in the last decade in pre-clinical studies due to their reliable expression of muscle wasting. However, these models succumb to tumor burden quickly, with overall survival less than 4 weeks and a cachexia window of only 2-3 weeks [10,11]. Additionally, these models do not express a common human cancer genotype or phenotype, particularly when implanted ectopically [30,31]. Others have addressed longevity using transgenic models, such as KPP and Apc^Min/+^, yielding important observations regarding longitudinal changes in cachexia [12,13]. However, these models express less common genotypes (i.e. mutations to *PTEN* and *APC*) compared to common human cancer from their organs of origin, limiting translational utility[32,33]. Conversely, Michaelis *et. al.* demonstrated that the KPC orthotopic model reliably expressed cachexia while retaining human pancreatic cancer genotype and phenotype [26]. Additionally, the KPC orthotopic model is syngeneic rather than transgenic or immunocompromised, which permits natural development of inflammatory biology; and this approach has successfully uncovered novel inflammatory mechanisms related to cachexia [34]. However, the limited survival window for the KPC orthotopic model prompted our need to extend its cachexia window using the LO approach. We hope this strategy will continue to be employed as a technique to develop translationally relevant cancer cachexia models.

Generation of this LO model facilitated time-based stratification of distinct physiologic events (Fig. 5C). Decline in gait speed occurred before even muscle fiber atrophy, suggesting that it occurs in the “pre-cachectic” phase. This mirrors Baltgalvis *et al.*, wherein voluntary wheeling running distance decreased prior to atrophy in the APC^Min/+^ model[13], while recent work by *Delfinis et al* also showed that muscle weakness precedes atrophy in C26 animals due to mitochondrial stress[35]. These combined findings suggest that early metabolic derangements lead to muscle fatigue, which is a precursor event for cachexia. Compared to changes in both skeletal and cardiac muscle bulk, we found that splenomegaly, metastases, and changes in anxiety are relatively late events, suggesting a distinct “refractory” phase in this model [3]. Intramuscular collagen deposition occurred as an intermediate event, leading to the intriguing idea that muscle fibrosis may represent the transition point from cachexia to refractory disease. Until now, the mechanisms that regulate transition to refractory cachexia have not been well investigated and the LO approach provides an opportunity to differentiate early and late events.

A major area of confusion in cancer-associated cachexia is the inability to distinguish factors that are specific to cancer progression versus factors that are unique to cachexia [36]. Functional outcome measures frequently used in the study of cancer patients are assumed to have relevance to the cachexia population but are not routinely compared to non-cachectic populations [37]. At the pre-clinical level, studies often ignore model heterogeneity or even stratify populations by tumor burden, rather than cachexia severity. We instead exploited the heterogeneity of the LO model to identify distinct cachectic and non-cachectic sub-populations via PCA (Fig. 5A). Using regression analysis to tie specific outcomes to cachexia, cancer, or both conditions, we found that gait speed was linked to both cachexia and cancer, while grip strength was only linked to cancer. Future studies can couple the LO approach with PCA to better identify factors that are specific to cachexia versus cancer.

Recent failures in cancer cachexia clinical trials have prompted many to suggest that a new approach is needed to developing cachexia specific outcome measures[3]. Clinical trials using anamorelin showed improvement in lean body mass but not hGS [38], leading to lack of approval by the FDA. The ultimate goal of using physical performance metrics is to objectively determine a patient’s physical functional ability and their quality of life. Recent studies suggest that direct measurement of functional independence may be promising for clinical trials with cachexia patients[2,39]. Yet, we demonstrate that change in hGS does not correlate with functional independence, similar to other recent studies showing a lack of sensitivity in hGS as a functional outcome in cachexia populations[6]. Because of this lack of sensitivity of hGS, alternative metrics have been proposed, including patient-reported outcome measures, timed-up-and-go, six-minute walk test, and actigraphy[37]. In this study, we show that change in 6MWT can act as a proxy measure for functional independence. This correlation held true in patients identified by each cachexia marker and was independent of age and disease recurrence status. These findings are consistent with prior studies in age-related sarcopenia suggesting that gait speed is an early indicator of clinical decline [40]. We speculate that since gait reflects the synchronized function of several muscle groups, it may be a more sensitive functional measure compared to handgrip, which involves only strength of distal muscles that change in mass relatively late in disease.

One limitation of our current approach is that gross muscle wasting below baseline is not detected for the first 4 weeks of the model, though fiber level atrophy is detected as early as 3 weeks post-implantation. We noted that control and tumor mice experienced muscle growth during those 4 weeks, decreasing the ability to sensitively detect simultaneous wasting. In post-hoc within cohort analysis, there is significant decline in gross muscle mass starting at 5 weeks, suggesting 5 weeks would be appropriate for future interventional studies directed at cachexia while interventions initiated at earlier time points may be more relevant to pre-cachexia. In future studies, we recommend advancing animal age to at least 20 weeks, when additional muscle growth is less likely. An additional limitation within this study is the IRF setting. IRF patients are a selected cohort that may not reflect the general cancer or cachexia population. Though we previously demonstrated a high prevalence of cachexia in our IRF cancer cohort [19] and a connection between functional independence and cachexia beyond the IRF setting [39]. Also, the median length of stay for inpatient rehabilitation does not permit collection of long-term outcomes, including weight recovery, thus we are unable to estimate if muscle bulk, which takes months to increase, also improved with function in this time frame. Future prospective and longitudinal studies that track cachexia patients through both inpatient and outpatient rehabilitation can further investigate the relationship between muscle mass and function recovery.

In summary, we propose that future pre-clinical studies of interventions intended to impact physical function should take a similar extended modeling approach and monitor gait speed as the primary outcome. Similarly, at the clinical level, clinical trials involving cachexia patients may be able to detect changes in function using 6MWT or similar gait related measures more sensitively than changes in grip strength.

## Author contributions

IR, AW, BBM, MBD, and RLL were involved with designing research studies, conducting experiments, acquiring data, analyzing data, and writing the manuscript. DS, TP, DM, AB, EM, DA, CF, RP and PJ were involved with conducting experiments, acquiring data, analyzing data, and editing the manuscript.

## Acknowledgements

We thank the Behavioral Phenotypic Core, Comprehensive Metabolic Core, and Center for Translational Imaging Core at Northwestern University. Model figures created with <u>BioRender.com</u>. This work was supported by a Foundational grant from the Catalyst Grant Program of the Shirley Ryan AbilityLab. IR is supported by career development award through the National Institute of Arthritis and Musculoskeletal and Skin Diseases (K08 AR081391). This work was also supported in part by Research Career Scientist Award Number IK6 RX003351 from the United States (U.S.) Department of Veterans Affairs Rehabilitation R&D (Rehab RD) Service.

## Ethical standards

Animal studies were approved by our Institutional Animal Care and Use Committee. The human subjects study was approved by our Institutional Review Board and received exempt status from consent procedures due to retrospective design.

