## Supplemental Methods, Figures, and Tables for "Gait speed is a biomarker of cancer-associated cachexia decline and recovery"

**Ex vivo tissue and histologic processing**

The tibialis anterior (TA) frozen samples were embedded in tragacanth gum, cut into three levels with 300 µm in between levels, and serial sectioned into three slides of 8 µm thickness, beginning at the proximal end of the muscle. Slides were then stained with laminin-GFP (Sigma, L9393) and DAPI counterstain. Tilescan images were obtained on a Leica CTR advanced microscope with the FLUO-green setting. Myovision parameters were set to the following- pixel scaling: 0.323, min area: 100, and max area: 9000. Fiber detection was carried out for each sample using a 20x tilescan of the second level from the proximal third of the laminin-stained TA. A minimum of 800 fibers were measured from each sample. For liver metastasis quantification, 3 sequential sections, 15 µm apart were made and then viewed to determine presence of metastases. Lungs were stained with H&E, and 5 sequential slides, 10 uM apart, were then viewed and metastases were counted. Final metastases values were determined by dividing the total number of metastases counted by the 5 slides. Spleens were stained for CD4 (abcam, ab183685). Tilescan image measurements were taken using a scale bar to “set scale” and calibrate ImageJ software. Next, the freehand selection function was used to outline the entire spleen perimeter and each germinal center (GC). The marginal zone (MZ) was calculated using 10 representative zones around GCs. All images were obtained using a LeicaCTR6500 microscope on the bright field (BF) setting.

**Biochemical analysis**

Enzyme linked immunoassays (ELISA) was performed on mouse serum using the Mouse Pentraxin2/SAP Quantikine ELISA kit (R&D Systems, MPTX20). Serum samples were diluted 1:4 to obtain results within the expected physiological levels and then processed according to manufacturer’s instructions. GDF15 was measured with a custom Mouse Luminex Discovery Assay (R&D Systems, LXSAMSM-05) following the manufacturer’s instructions. The assay was run with the help of the Comprehensive Metabolic Core at Northwestern University.

**CT and MRI imaging**

CT images were obtained using the IVIS SpectrumCT (Perkin Elmer, Waltham, MA) with animals in the supine position. Muscle volume was approximated using the index of muscle mass technique applied at the distal hindlimb as previously described [1]. MRI images were obtained in the Center for Translational Imaging at Northwestern University on the Bruker 7T ClinScan MRI System (Billerica, MA). ITK-Snap ([www.itksnap.org](http://www.itksnap.org)) was used to process MRI Double Echo Steady State sequences. The most lateral sagittal plane sequence where both the tibia and femur were in plane was identified. Quadriceps volume was then measured from above the patella.

**Animal behavior analysis**

Mouse sepsis score was used to track sepsis like behavior [2]. Individual limb grip strength was performed as previously described [3,4]. Once the animal was stabilized, the meter was zeroed, and the restrained mouse was allowed to grasp the bar with one limb while the researcher slowly pulled the animal back. Maximum grip strength out of 10 trials for each individual limb was recorded by the meter at the time the mouse released the bar. For rotarod, mice were subjected to four, 5-minute intervals on the rotarod drum rotating at a constant speed of 12 RPM (RotaRod 3375-5, TSE Systems, Bad Homburg, Germany). The length of time each animal was able to maintain its balance on the drum was measured [5].

**Human functional outcome measures**

Secondary outcomes included lack of independence at discharge based on discharge setting (return to acute care or admission to a skilled nursing facility/long term acute care facility) or need for home health services due to inability to exit their home for outpatient medical care.

**Principal component analysis of animal data**

Principal component analysis (PCA) was performed by Graphpad Prism 9.0. Variables included in the PCA were quadriceps mass, cardiac mass, primary tumor mass, peritoneal metastasis presence, and splenomegaly presence. Principal component linear regression analysis was performed to analyze the relationship between variables within the PCA and functional outcome measures. To observe whether there were any trends of time since implantation for any of the PCA components, we plotted the data from the PCA analysis integrating in days since injection information. To do so, we plotted the data points in 3-D (x – PC1, y – PC2, z – days since inject) and then overlaid a surface mesh that was interpolated across data points using the MATLAB function *griddata (*The MathWorks Inc., Natick, MA v2022a). We then created a 3-d vector plot following the contours of the surface mesh towards increasing time since injection using the MATLAB function *gradient (*The MathWorks Inc., Natick, MA v2022a).

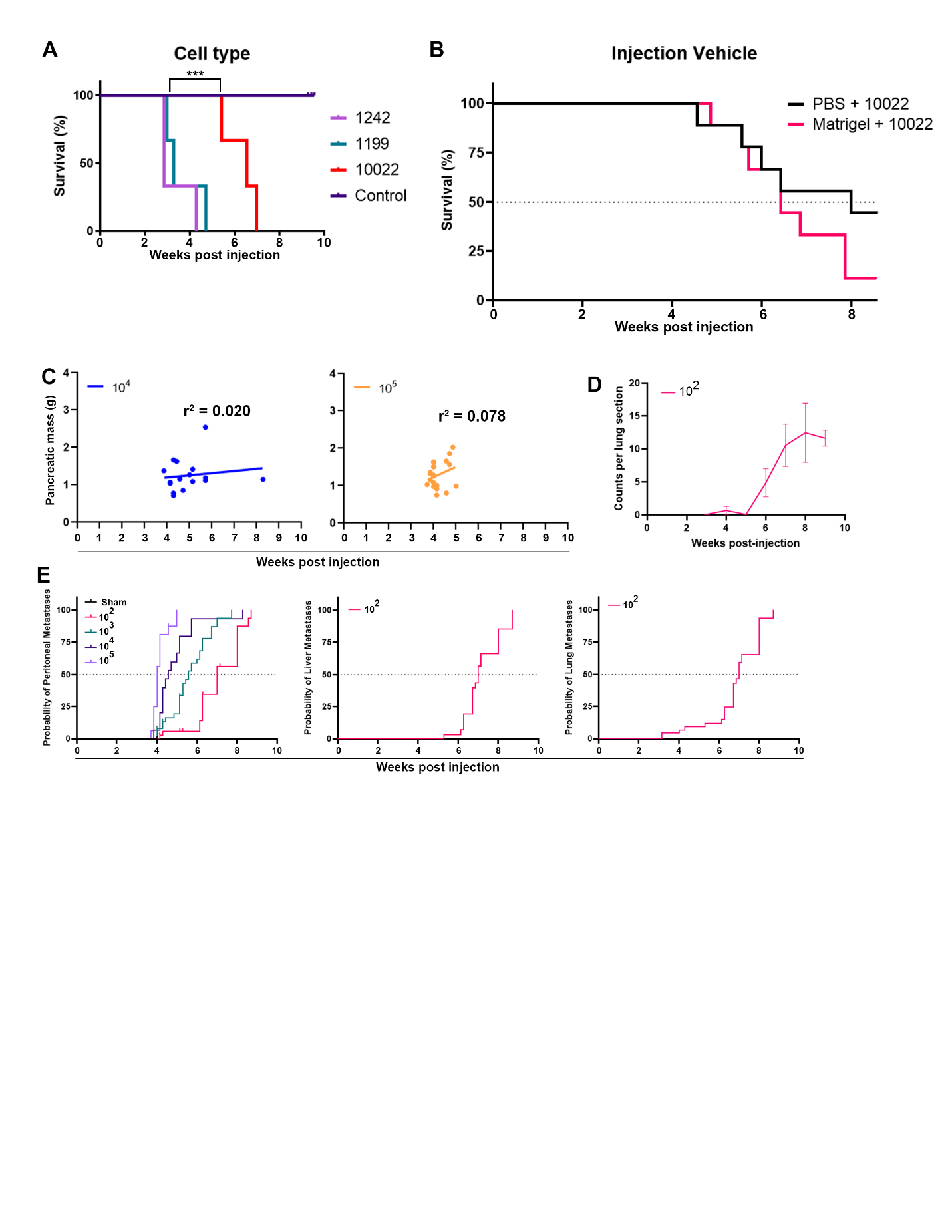
**Supplementary Fig. 1. Extending KPC orthotopic model through manipulation of cell type and injection vehicle.** (A) Kaplan-Meier survival analysis of cohorts implanted with 10000 cells of distinct KPC cell lines: 1242, 1199, and 10022 compared to sham control. n=3 per cohort. * indicates p < 0.05. (B) Kaplan-Meier survival analysis of cohorts implanted with 100 cells mixed with either sterile PBS of 1:4 dilution of Matrigel. (C) Simple linear regression of *ex vivo* pancreatic/primary tumor weight vs. days post implantation at Sham, 10000, and 100000 doses. (D) Counts of lung metastases per H&E tissue section in low dose model over time in serial cohorts. (E) Kaplan-Meier curves for peritoneal, liver, and lung metastases over time at varying doses of 10022 cells. Median time to metastasis in all three anatomic locations was 7 weeks.

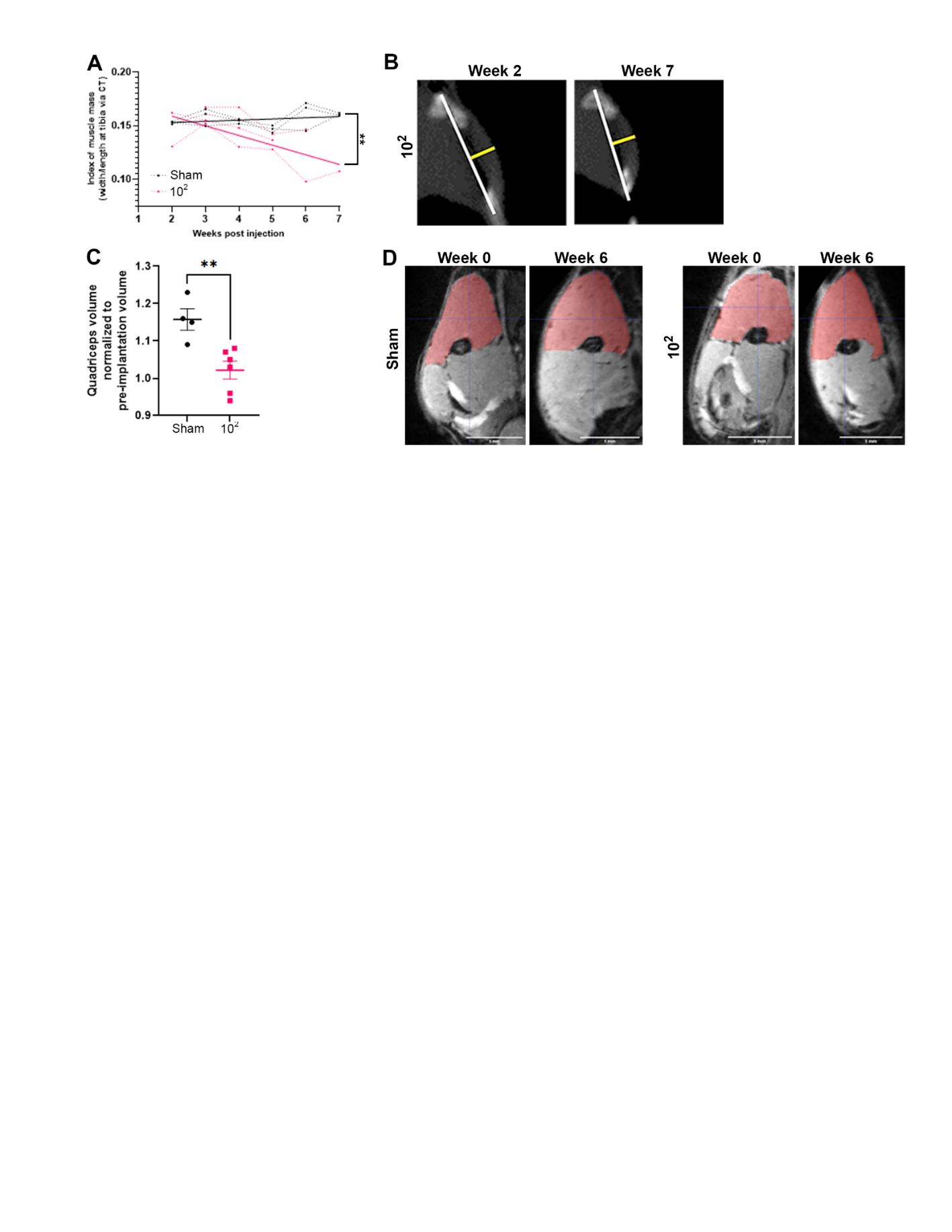
**Supplementary Fig. 2. Imaging of muscle wasting in low dose KPC orthotopic model.** (A-B) Index of muscle mass in Sham and 100 KPC cell implanted animals as repeated measure over time along with representative images showing muscle thickness and midpoint of tibia. n=3 per cohort. * indicates p < 0.05 for difference in linear regression calculated slope between cohorts. (C-D) Quadriceps volume measured via MRI over time in Sham and 100 KPC cell implanted animals along with representative images showing measured area at midpoint of femur. In a two-way ANOVA, “a” indicates p < 0.05 for time factor, “b” indicates p < 0.05 for cell dose factor, and “c’” indicates p < 0.05 for interaction between time and cell dose. “**” indicates p < 0.01 in unpaired t-test.

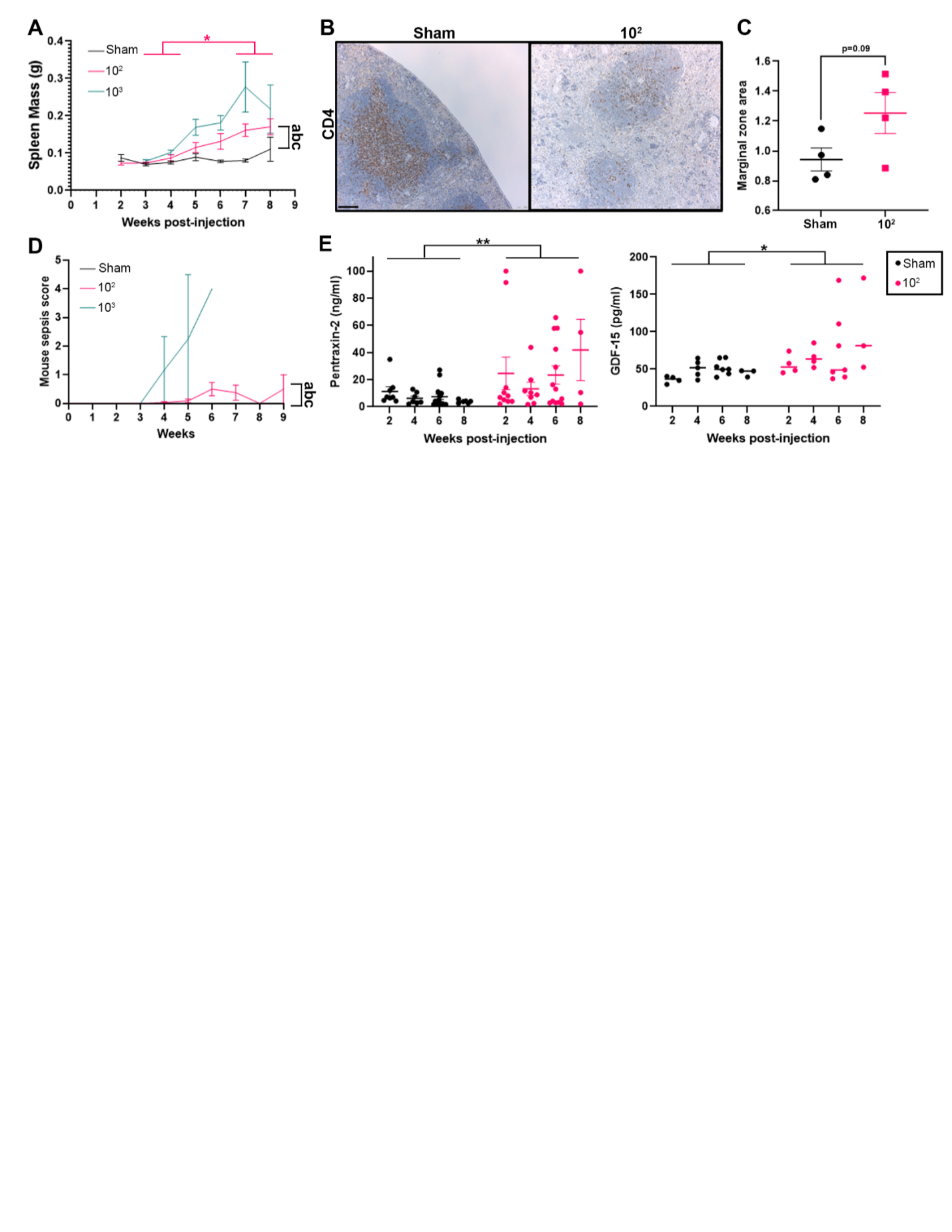
**Supplementary Fig. 3. Systemic inflammation over time in low dose KPC orthotopic model.** (A) *Ex vivo* spleen mass over serial weekly cohorts in SHAM, 100, and 1000 implanted animals. (B) Representative CD4 stained images of spleens from SHAM and 100 cohorts. Size bar = 100 µm. (C) Total area of marginal zones per section in SHAM and 100 spleens. (D) Mouse sepsis score as repeated measure over time in SHAM, 100, and 1000 implanted mice. (E) Serum pentraxin and GDF-15 levels at weeks 2, 4, 6, and 8 post implantation in SHAM and 100 mice. For panels A, n = 7-12 per weekly cohort per cell dose, other than week 9 (n=5 per cell dose). For panel D, n = 31-37 for SHAM and 100 cohorts and n = 8 for 1000 cohort. For panel E, n= 4-8 per time point for Pentraxin-2 and n= 3-7 per time point for GDF-15. In a two-way ANOVA, “a” indicates p < 0.05 for time factor, “b” indicates p < 0.05 for cell dose factor, and “c’” indicates p < 0.05 for interaction between time and cell dose. Pink * indicates p < 0.05 in Tukey post-hoc analysis between time points within the 100 cohort. Black * indicates p < 0.05 for tumor status in a two-way ANOVA.

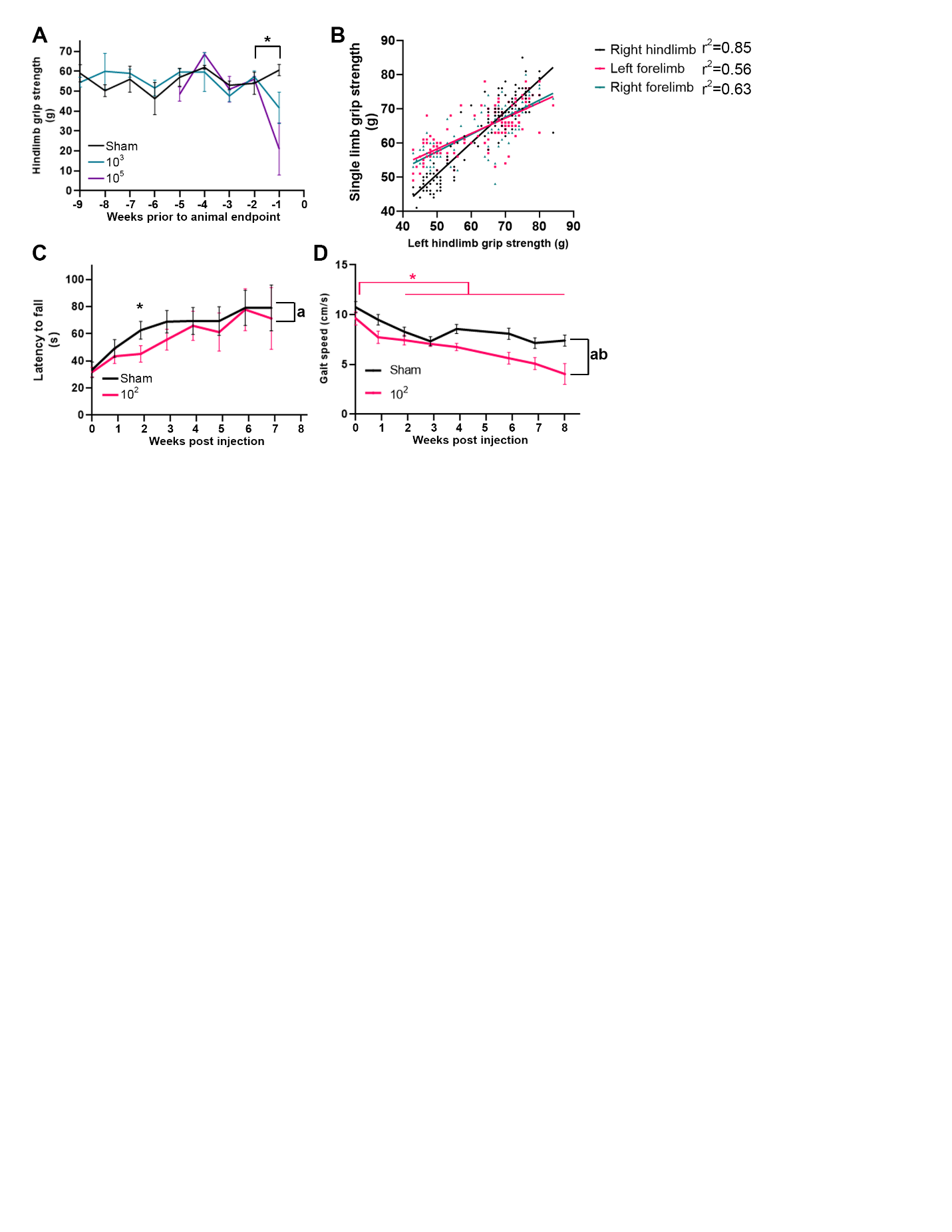

**Supplementary Fig. 4. Grip strength and rotarod in KPC orthotopic model.** (A) Repeated measure hindlimb grip strength in Sham, 1000, and 100000 implanted animals vs. days prior to animal endpoint (determined by institutional criteria). n = 4 per cohort. * indicates p < 0.05 in within cohort analysis of tumor implanted animals. (B) Correlation between grip strength of individual limbs with corresponding r^2^ values in 100-cell dose implanted animals. (C) Repeated measure latency to falling from rotarod apparatus of Sham and 100 cohorts, * indicates p < 0.05 between cohort analysis at that select time point. (D) Gait speed in two dimensional tracking between sham and 100 cohorts in open field maze. n = 26-33 per cohort; in a two-way ANOVA, “a” indicates p < 0.05 for time factor, “b” indicates p < 0.05 for cell dose factor, and “c’” indicates p < 0.05 for interaction between time and cell dose. Pink * indicates p < 0.05 in Tukey post-hoc analysis between time points within the 1e2 cohort.

**Supplementary Table 1. Principal Component Analysis Calculations**

| **PC summary** | **PC1** | **PC2** | **PC3** | **PC4** | **PC5** |  |
| --- | --- | --- | --- | --- | --- | --- |
| **Eigenvalue** | 2.4 | 1.5 | 0.48 | 0.36 | 0.21 |  |
| **Proportion of variance** | 48.53% | 30.24% | 9.67% | 7.29% | 4.27% |  |
| **Cumulative proportion of variance** | 48.53% | 78.77% | 88.44% | 95.73% | 100.00% |  |
| **Component selection** | Selected | Selected |  |  |  |  |
|  | **Loadings** | | **Contributions of variables** | | **Eigenvectors** | |
|  | **PC1** | **PC2** | **PC1** | **PC2** | **PC1** | **PC2** |
| **quadriceps** | -0.52639 | 0.754556 | 0.114204 | 0.376522 | -0.33794 | 0.613614 |
| **pancreas** | 0.820133 | 0.298131 | 0.277221 | 0.058779 | 0.526518 | 0.242443 |
| **heart** | -0.47358 | 0.794094 | 0.092435 | 0.417015 | -0.30403 | 0.645767 |
| **spleen** | 0.821016 | 0.352281 | 0.277819 | 0.08207 | 0.527085 | 0.286479 |
| **peritoneal metastasis** | 0.760418 | 0.314989 | 0.238321 | 0.065614 | 0.488182 | 0.256152 |

**Supplementary Table 2. Demographics of cancer patients receiving inpatient rehabilitation with grip strength and 6MWT measured pre- and post- rehabilitation.**

| WLGS | 0 | 1 | 2 | 3 | 4 | p-value^1^ |
| --- | --- | --- | --- | --- | --- | --- |
| n | 35 | 24 | 26 | 44 | 27 |  |
| Age (median) | 68 | 57 | 63 | 60 | 71 | 0.0021 |
| Gender (% male) | 46% | 29% | 50% | 52% | 67% | 0.1121 |
| Cancer (n) | |  |  |  |  | 0.0949 |
| Breast | 7 | 2 | 3 |  | 1 |  |
| GI/Pancreas/HPB | 1 | 1 | 1 | 3 | 4 |  |
| GU | 5 | 1 | 3 | 1 | 1 |  |
| gyn |  |  | 1 |  |  |  |
| H&N |  |  | 1 | 1 |  |  |
| Hematologic | 4 | 4 | 2 | 12 | 9 |  |
| lung | 1 |  | 2 | 4 | 5 |  |
| MSK |  | 1 |  | 1 |  |  |
| Other | 1 |  | 1 |  |  |  |
| Primary intracranial | 15 | 12 | 11 | 19 | 6 |  |
| Primary Spine | 1 | 2 | 1 | 1 |  |  |
| Skin |  | 1 |  | 2 | 1 |  |
| Recurrent cancer | 18 | 13 | 19 | 29 | 22 | 0.0906 |

^1^Chi-squared analysis.

**Supplementary Table 3. Linear regression between FIM motor, 6MWT, and grip strength gains in cancer patients with muscle wasting**

|  | Unstandardized Coeff | SE | 95% CI | R-squared | p-value^1^ |
| --- | --- | --- | --- | --- | --- |
| 6MWT change vs Motor FIM Gain | | |  |  |  |
| WLGS = 0 | 3.49 | 1.06 | 1.342 to 5.643 | 0.249 | 0.002 |
| WLGS = 1 | 4.52 | 1.46 | 1.498 to 7.541 | 0.304 | 0.005 |
| WLGS = 2 | 2.63 | 1.83 | -1.137 to 6.405 | 0.080 | 0.162 |
| WLGS = 3 | 3.88 | 1.47 | 0.9185 to 6.845 | 0.143 | 0.012 |
| WLGS = 4 | 4.40 | 1.50 | 1.317 to 7.489 | 0.257 | 0.007 |
| Fearon et al criteria | 3.92 | 0.90 | 2.134 to 5.696 | 0.178 | <0.0001 |
| PNI < 40 | 3.89 | 0.93 | 2.015 to 5.766 | 0.274 | <0.0001 |
| GS change vs Motor FIM Gain | |  |  |  |  |
| WLGS = 0 | 0.377 | 0.13 | 0.1116 to 0.6420 | 0.283 | 0.008 |
| WLGS = 1 | -0.346 | 0.24 | -0.8836 to 0.1912 | 0.171 | 0.182 |
| WLGS = 2 | -0.089 | 0.10 | -0.3017 to 0.1237 | 0.039 | 0.392 |
| WLGS = 3 | 0.067 | 0.15 | -0.2294 to 0.3640 | 0.007 | 0.646 |
| WLGS = 4 | 0.257 | 0.17 | -0.1036 to 0.6175 | 0.090 | 0.154 |
| Fearon et al criteria | 0.039 | 0.09 | -0.1404 to 0.2175 | 0.003 | 0.669 |
| PNI < 40 | 0.234 | 0.13 | -0.0229 to 0.4912 | 0.118 | 0.07 |

^1^Univariate linear regression.

**Supplementary Table 4. Linear regression between FIM motor, 6MWT, and grip strength gains in male and female cancer patients using Fearon et al cachexia**

|  | Unstandardized Coeff | SE | 95% CI | R-squared | p-value^1^ |
| --- | --- | --- | --- | --- | --- |
| 6MWT change vs Motor FIM Gain - males | | |  |  |  |
| Cachexia | 3.989 | 1.217 | 1.543 to 6.434 | 0.1767 | 0.0019 |
| No cachexia | 3.038 | 1.533 | -0.1268 to 6.203 | 0.1406 | 0.0591 |
| 6MWT change vs Motor FIM Gain - females | | | | | |
| Cachexia | 3.947 | 1.408 | 1.091 to 6.802 | 0.1792 | 0.0081 |
| No cachexia | 3.594 | 1.115 | 1.341 to 5.848 | 0.2063 | 0.0025 |
| GS change vs Motor FIM Gain - males | | | | | |
| Cachexia | 0.1868 | 0.1517 | -0.1297 to 0.5033 | 0.07045 | 0.2326 |
| No cachexia | 0.7287 | 0.3202 | -0.05484 to 1.512 | 0.4633 | 0.0632 |
| GS change vs Motor FIM Gain - females | | | | | |
| Cachexia | 0.01919 | 0.2034 | -0.4119 to 0.4503 | 0.0005561 | 0.9260 |
| No cachexia | 0.1931 | 0.1982 | -0.2272 to 0.6133 | 0.05598 | 0.3445 |

^1^Univariate linear regression.

**Supplementary Table 5. Linear regression between FIM cognitive, 6MWT, and grip strength gains in cancer patients with muscle wasting**

|  | Unstandardized Coeff | SE | 95% CI | R-squared | p-value^1^ |
| --- | --- | --- | --- | --- | --- |
| 6MWT change vs Cognitive FIM Gain | | |  |  |  |
| WLGS = 0 | 1.00 | 4.37 | -7.885 to 9.889 | 0.0016 | 0.820 |
| WLGS = 1 | 0.48 | 4.14 | -8.108 to 9.071 | 0.0006 | 0.909 |
| WLGS = 2 | 1.57 | 5.68 | -10.16 to 13.29 | 0.0032 | 0.785 |
| WLGS = 3 | 4.99 | 3.08 | -1.225 to 11.20 | 0.0588 | 0.113 |
| WLGS = 4 | 5.71 | 3.78 | -2.061 to 13.49 | 0.0839 | 0.143 |
| Fearon et al criteria | 5.13 | 2.19 | 0.7680 to 9.484 | 0.0585 | 0.022 |
| PNI < 40 | 6.45 | 2.65 | 1.117 to 11.78 | 0.1141 | 0.019 |
| GS change vs Cognitive FIM Gain | |  |  |  |  |
| WLGS = 0 | 0.5047 | 0.223 | 0.00790 to 1.002 | 0.3388 | 0.0471 |
| WLGS = 1 | -0.1693 | 0.3094 | -1.028 to 0.6898 | 0.06966 | 0.6133 |
| WLGS = 2 | 0.1838 | 0.1415 | -0.1277 to 0.4952 | 0.1329 | 0.2206 |
| WLGS = 3 | 0.5069 | 0.2935 | -0.1188 to 1.133 | 0.1658 | 0.1047 |
| WLGS = 4 | 0.07577 | 0.4026 | -0.7939 to 0.9454 | 0.002718 | 0.8536 |
| Fearon et al criteria | -0.1951 | 0.2095 | -0.6132 to 0.2230 | 0.01278 | 0.355 |
| PNI < 40 | 0.05278 | 0.5989 | -1.252 to 1.358 | 0.0006467 | 0.9312 |

^1^Univariate linear regression.

**Supplementary Table 6. Univariate and multivariate logistic regression analysis of level of independence on discharge from rehabilitation in cancer patients with Weight Loss Grading Scale > 0**

| Discharge Independence |  | OR | 95% CI | p-value^1^ |
| --- | --- | --- | --- | --- |
| Univariate | 6MWT gain | 0.9947 | 0.9895 to 0.9993 | 0.0316 |
|  | GS gain | 1.015 | 0.9603 to 1.074 | 0.6071 |
| Multivariate | 6MWT gain | 0.9929 | 0.9841 to 1.000 | 0.081 |
|  | GS Gain | 1.031 | 0.9529 to 1.122 | 0.4436 |
|  | Age | 1.034 | 0.9860 to 1.090 | 0.1826 |
|  | Recurrent disease | 3.373 | 0.7816 to 18.40 | 0.1215 |

^1^Univariate and multivariate linear regression.
